## Supplementary Table 2 for "Zinc finger homeobox-3 (ZFHX3) orchestrates genome-wide daily gene expression in the suprachiasmatic nucleus"

Supplementary Table 2: Circadian wheel running analysis comparing *Zfhx3*^Flox/Flox^;UBC animals dosed with Tamoxifen (+Tamoxifen) to those without Tamoxifen dosing (-Tamoxifen).

|  |  | Circadian period (Tau) (hours) | Amplitude (PN value) | Intraday variability (IV) | Interdaily stability (IS) | Activity onset in LD (hours) |
| --- | --- | --- | --- | --- | --- | --- |
| Female | -Tamoxifen | 23.68 ±0.03 | 3375 ±643 | 0.687 ±0.103 | 0.255 ±0.036 | 12.03 ±0.033 |
|  | +Tamoxifen | 23.22 ±0.12 | 1809 ±674 | 0.824 ±0.105 | 0.059 ±0.013 | 11.48 ±0.202 |
| Male | -Tamoxifen | 23.74 ±0.04 | 2514 ±773 | 0.929 ±0.185 | 0.28 ±0.039 | 12.04 ±0.031 |
|  | +Tamoxifen | 23.33 ±0.18 | 140 ±71 | 1.695 ±0.119 | 0.042 ±0.005 | 10.21 ±0.707 |
| Repeat Measures ANOVA analysis | Tamoxifen Dosing | F(1,26)=18.39; **p=0.0002** | F(1,26)=9.05; **p=0.0058** | F(1,26)=11.05; **p=0.0026** | F(1,26)=53; **p<0.0001** | F(1,26)=15.71; **p=0.0005** |
|  | Sex | F(1,26)=0.66; p=0.4223 | F(1,26)=3.7; p=0.0642 | F(1,26)=16.78; p=0.0004 | F(1,26)=0.02; p=0.8873 | F(1,26)=4.332; p=0.0474 |
|  | Tamoxifen Dosing X Sex | F(1,26)=0.053; p=0.8197 | F(1,26)=0.38; p=0.5422 | F(1,26)=5.36; p=0.0287 | F(1,26)=0.53; p=0.4731 | F(1,26)=4.516; p=0.0432 |
